# A universal stress protein acts as a metabolic rheostat controlling carbon flux in mycobacteria

**DOI:** 10.1101/2022.06.10.495724

**Authors:** Kathryn E. Lougheed, Michael Thomson, Lukasz S. Koziej, Gerald Larrouy-Maumus, Huw D Williams

**Affiliations:** Department of Life Sciences, Imperial College London, London, United Kingdom; MRC Centre for Molecular Bacteriology and Infection, Imperial College London, London, United Kingdom

## Abstract

Universal stress proteins (USPs) are ubiquitous amongst prokaryotes and have empirically determined roles in adaptation to stress, while their specific biochemical function in cellular processes is opaque. The most frequently encountered USP is a small ~15kDa protein comprised of a single domain (PF00582). In *Mycobacterium tuberculosis* Rv1636 is the sole single domain protein amongst its 10 USPs. We were unable to delete rv1636 from *M. tuberculosis*, supporting previous data that it is essential for growth. We deleted its orthologue, MS3811, from *M. smegmatis*, and in exponential phase competition experiments we observed that the advantage switched between wildtype and the mutant depending on the growth environment. In rich medium the wildtype had the competitive advantage, but in a defined medium with a single carbon source the competitive advantage switched, with the mutant rapidly taking over the culture. We hypothesised the USP is regulating metabolic flux, with its deletion leading to increased flux and more rapid turn-over of central-carbon metabolites. We tested this by performing ^13^C-stable isotope tracing, using [U-^13^C_6_] glucose as sole carbon source for growth of the parental and deletion strains. The findings were unequivocal; deletion of the USP led to an increase in label incorporation in central carbon metabolites and a profound change in the isotopologue distribution. Furthermore, *in vivo* protein-crosslinking provided evidence that Rv1636 interacts directly with key central metabolic enzymes, including the glycolytic enzymes pyruvate kinase, pyruvate dehydrogenase, pyruvate synthase. We propose the mycobacterial single domain USP acts as a metabolic rheostat to regulate central carbon metabolism.

**Importance:** There are more than 62,000 universal stress proteins (USPs) found mostly in prokaryotes, but very few examples where the role of a USP is unequivocally linked to a specific biological or biochemical function. For the first time we were able to assign a role in metabolic regulation to a bacterial single domain universal stress protein. We demonstrate a mycobacterial USP regulates flux through central catabolic pathways, and our evidence, based on ^13^C-stable isotope tracing, indicates that the USP acts as a brake on carbon flux, and we assign to it a role as a metabolic rheostat.

## Introduction

Universal stress proteins (USP_s_) are ubiquitously found in bacteria, archaea, plants and some invertebrates and have regulatory and protective roles in adapting organisms to external stresses (1–3).

Universal stress proteins (USP_s_) are characterised by containing at least one USP domain (PF00582), which comprises an α/β core structure consisting of 140 to 160 conserved amino acid residues forming five β-strands and four α-helices. It can be associated with a variety of other functional motifs, but most USP_s_ comprise either a single USP domain or two tandem USP domains (1, 3, 4). Of 62,462 USP_s_ in the PFAM database from more than 7,200 species, 33,564 are single domain USP_s_ and more than 11,000 are tandem domain proteins (http://pfam.xfam.org/). USP domains are sub-divided into two groups. The UspEG-type show homology to the structure of the USP domain in the *Methanococcus jannaschi* protein MJ0577 (5) and contain a conserved ATP-binding motif in the C-terminal region [G-2X-G-9X-(S/T)], while those in the UspA-like group show homology to the UspA from *Haemophilus influenzae* and do not have an ATP-binding motif (6).

Most bacteria have multiple paralogs of USP_s_, but there is large variation in the number present in a particular bacterial species. This is illustrated in the actinobacteria; *Mycobacterium leprae* has 1, *M. tuberculosis* 10, *M. smegmatis* 17 and some *Streptomyces* species more than 30 different USP_s_ proteins encoded in their genome (7).

In bacteria there is a consistent link between USP_s_ and survival under growth-arrested conditions, with adaptation to nutrient stress or energy stress a common theme. Universal stress protein A (UspA) was originally discovered in *Escherichia coli* as a protein that was overexpressed during environmental stress that led to growth arrest (8), where it enhances cell survival and it and related proteins have been described as having a general “stress endurance” activity (6, 9). After this discovery Usp-type proteins were identified in most species of bacteria, archaea, plants and simple metazoans, but not vertebrates (1, 4).

An *E. coli uspA* mutant showed premature death in stationary phase (8–10), while a *Salmonella typhimurium uspA* mutant survived carbon or phosphorous starvation poorly and had increased sensitivity to oxidative stress (11). In *Pseudomonas aeruginosa* USP_s_ supported its survival in anaerobic stationary phase and in the presence of a respiratory uncoupler (12, 13). USP_s_ are also involved in the switch to anaerobiosis in *Salmonella enterica* serovar Typhimurium (14), while in the periodontal pathogen *Porphyromonas gingivalis* they have a role in anaerobic biofilm survival (15). In the photosynthetic bacterium *Rhodobacter sphaeroides* energy stress, in the form low light intensity, upregulates Usp protein expression (16). However, despite a body of evidence linking them to diverse cellular processes only few studies have successfully investigated the biochemical roles of USP_s_ and mechanistic insights into their function have proved elusive.

*M. tuberculosis* is an obligate aerobe that upregulates multiple USP_s_ when it becomes energy starved due to a switch to hypoxic growth conditions prior to entry into a non-replicating persistent state (7, 17). The genome of *M. tuberculosis* encodes for 10 USP proteins. One is a single domain USP and eight are tandem domain proteins and all have a putative ATP-binding motif (7). The final protein is the sensor kinase KdpD that regulates the high affinity K^+^-transport system KdpFABC in response to K^+^-limitation or salt stress and contains a USP domain (18). Some *M. tuberculosis* USP_s_ are upregulated by stresses that include low pH, nitric oxide, UV light and mitomycin C and they are expressed in macrophages and the expression of six tandem domain USP_s_ are regulated by the dormancy regulator DosR in response to hypoxia (19) (7) (20). However, mutants in four of these (Rv1996, Rv2005c, Rv2026c and Rv2028c) lead to no discernible phenotype in response to a wide variety of stresses and growth conditions (21). In contrast deletion of the USP gene *rv2623* led to an alteration of *M. tuberculosis* survival and persistence in macrophages and *rv2623* was highly upregulated under hypoxia and nitrosative stress (20). Recently it has been shown that Rv2623 is involved in the regulation of mycobacterial growth by its interaction with the ABC transporter Rv1747, suggesting a role of this USP in the metabolic activities of *M. tuberculosis* (22).

Rv1636 is the sole single domain USP in *M. tuberculosis* and it is ubiquitous and highly conserved in mycobacteria. It has a homologue MS3811 in *M. smegmatis* and its homologue is the only USP present in the obligate intracellular pathogen *M. leprae*. Intriguingly, Rv1636 and MS3811 can bind the second messenger cAMP (23). Our aim was to investigate the physiological role of this conserved single domain USP by using a combination of microbiology, molecular biology and metabolomics coupled to stable isotope tracing.

## Results

Given the stress response phenotypes reported for *usp* mutants from a range of bacteria we tested whether the survival of *M. smegmatis* Δ3811 was compromised under a range of stress conditions that included: cell wall stress (SDS up to 1% w/v) H_2_O_2_ (5mM), starvation in PBS, salt stress (5 M NaCl), ethanol exposure (5-10% v/v), heat shock (50°C for 7 h, 55°C for 5 h), and we determined antibiotic MICs for wildtype and the Δ3811 mutant (gentamycin, kanamycin, spectinomycin, rifampicin, INH, ethionamide, ofloxacin). No reproducible mutant phenotype was observed with any of these conditions and, furthermore, the *M. smegmatis* Δ3811 mutant showed no stationary phase-survival defect (data not shown). We measured the intracellular concentration of MS3811 in *M. smegmatis* using Multiple Reaction Monitoring Mass spectrometry (MRM-MS) and determined that it was about 50 nM in both exponential and stationary phase cells, indicated that this USP is constitutively produced (Fig. S1). This is contrast to the finding that many bacterial USP_s_ are expressed in response to growth arrest, including six *Mtb* USP_s_, (but not Rv1636) and many of those in *M. smegmatis*, which are induced in response to hypoxic stationary phase (19, 21, 24, 37) (1).

### The competitive advantage switches between wildtype and the single domain *usp* mutant depending on the growth environment

The constitutive expression of MSMEG_3811 suggests its role, unlike that of many other USP_s_, is not restricted to growth arrested cells and that it has a function in growing bacteria. We investigated this further using a sensitive competition assay. Equal numbers of *M. smegmatis* wildtype and ΔMSMEG_3811 mutant bacteria were inoculated into growth media at a density of about 5 x 10^7^ cfu ml^-1^ and the cultures maintained in exponential phase by daily sub-culturing into fresh medium. Cultures were plated onto medium without and with hygromycin to give the total bacterial count and that of the mutant (Hyg^R^), respectively.

In LB-Tween rich medium, which has multiple carbon sources, the wildtype has a clear competitive advantage over the ΔMSMEG_3811 mutant during exponential phase leading to it comprising more than 90% of the population after 14 days. However, the competitive advantage switches to the ΔMSMEG_3811 mutant when the strains are competed in a medium with a single carbon source. During competition in HdB medium with glucose as the carbon source the ΔMSMEG_3811 mutant rapidly takes over of the culture, and a similar outcome is seen when pyruvate, glycerol, acetate or cholesterol are the single carbon sources in the medium. This striking finding indicates that the loss of the MSMEG_3811 is advantageous with a single carbon source but a disadvantageous when multiple carbon sources are present, suggesting that MSMEG_3811, (and its homologues in *Mtb* and *M. leprae*) may have a regulatory role in coordinating metabolism.

### Deletion of MS3811 leads to an increase in the turn-over of central-carbon metabolites

The carbon-source dependent competition phenotypes led us to hypothesise that the conserved mycobacterial single domain USP regulates and optimises metabolic flux through central carbon metabolism.

To test the model that the USP modulate central carbon metabolism in *M. smegmatis*, we performed ^13^C-stable isotope tracing to investigate alteration in the turn-over of central carbon metabolites following growth in HdB medium supplemented with [U-^13^C_6_] glucose in the parental and deletion strains. As seen in Table 1, deletion of MSMEG_3811 leads to a significant increase in label incorporation in metabolites involved in central carbon metabolism. For example, compared to the WT parental strain, the ΔMSMEG_3811 mutant a 1.15-fold increase in total ^13^C incorporation is observed in pyruvate, 1.3-fold in L-alanine, 1.3-fold in succinic acid, 1.2-fold in oxoglutaric acid and 1.5-fold in citric acid.

**Table 1:** Percentage of ^13^C Carbon incorporation in a cross section of central carbon, nitrogen and amino acid metabolites in both WT parental and ΔMS3811 strains.

| Compound Name | %labelled wild-type | %labelled $\Delta\text{MS3811}$ | *p-value* |
| --- | --- | --- | --- |
| Pyruvic acid | 80.0 ± 1.9 | 92.5 ± 2.3 | 1.87E-05 |
| L-Alanine | 59.7 ± 3.7 | 76.8 ± 5.9 | 9.30E-04 |
| γ-Aminobutyric acid | 76.8 ± 0.9 | 83.3 ± 6.7 | 9.46E-02 |
| L-Serine | 68 ± 2.1 | 82.6 ± 6.9 | 2.05E-03 |
| L-Proline | 12.6 ± 3.8 | 53.8 ± 9.1 | 2.98E-05 |
| L-Valine | 45.8 ± 4.9 | 71.0 ± 10.8 | 2.49E-03 |
| Succinic acid | 50.0 ± 4.3 | 65.6 ± 6.6 | 3.34E-03 |
| L-Threonine | 47.4 ± 2.7 | 70.0 ± 6.9 | 2.78E-04 |
| L-Leucine | 5.7 ± 3.8 | 48.7 ± 14.8 | 5.17E-04 |
| L-Ornithine | 68.3 ± 3.0 | 74.7 ± 3.8 | 2.34E-02 |
| L-Aspartic Acid | 65.2 ± 2.2 | 72.1 ± 10.1 | 2.20E-01 |
| Oxoglutaric acid | 75.8 ± 0.8 | 88.4 ± 3.9 | 2.63E-04 |
| Glutamine | 72.8 ± 1.5 | 89.5 ± 3.1 | 9.95E-06 |
| L-Glutamate | 77.0 ± 0.9 | 84.0 ± 7.6 | 1.10E-01 |
| trans-Aconitate | 60.1 ± 7.6 | 80.8 ± 6.1 | 1.43E-03 |
| Citrulline | 87.9 ± 0.8 | 90.5 ± 2.4 | 7.49E-02 |
| Citric acid | 62.0 ± 0.9 | 90.1 ± 2.4 | 1.71E-08 |

Interestingly, even though for some metabolites the overall percentage of ^13^C in the metabolites does not change significatively, a profound change can be observed in the isotopologue distribution in the strain lacking MSMEG_3811. As is shown in Figure 2, the absence of MSMEG_3811 leads to an increase in labelling with isotopologues containing a high number of carbons and a decrease in the labelling with isotopologues containing a low number. For example, compared to the parental strain, in the ΔMSMEG_3811 strain, there was a 0.3-fold increase in M+3 in pyruvic acid. A similar pattern is seen in other metabolites such as citric acid – a 2-fold and 4-fold increase in M+5 and M+6 concomitant with a decrease in M+1 and M+2 isotopologues; L-glutamic acid – a 2-fold increase in M+5 concomitant with a decrease in M+1; trans-aconitate – a 2.5-fold increase in M+6 concomitant with a decrease in M+0; oxoglutaric acid – a 2.1-fold increase in M+5 concomitant with a 2.2-fold decrease in M+0; aspartic acid – a 1.9-fold increase in M+4 concomitant with a 1.8-fold decrease in M+1 and 1.3-fold decrease of M+2; succinic acid – a 1.25-fold and 2-fold increase in M+3 and M+4 concomitant with a 1.5-fold and 1.42-fold decrease in M+0 and M+1.

**Figure. 1.**
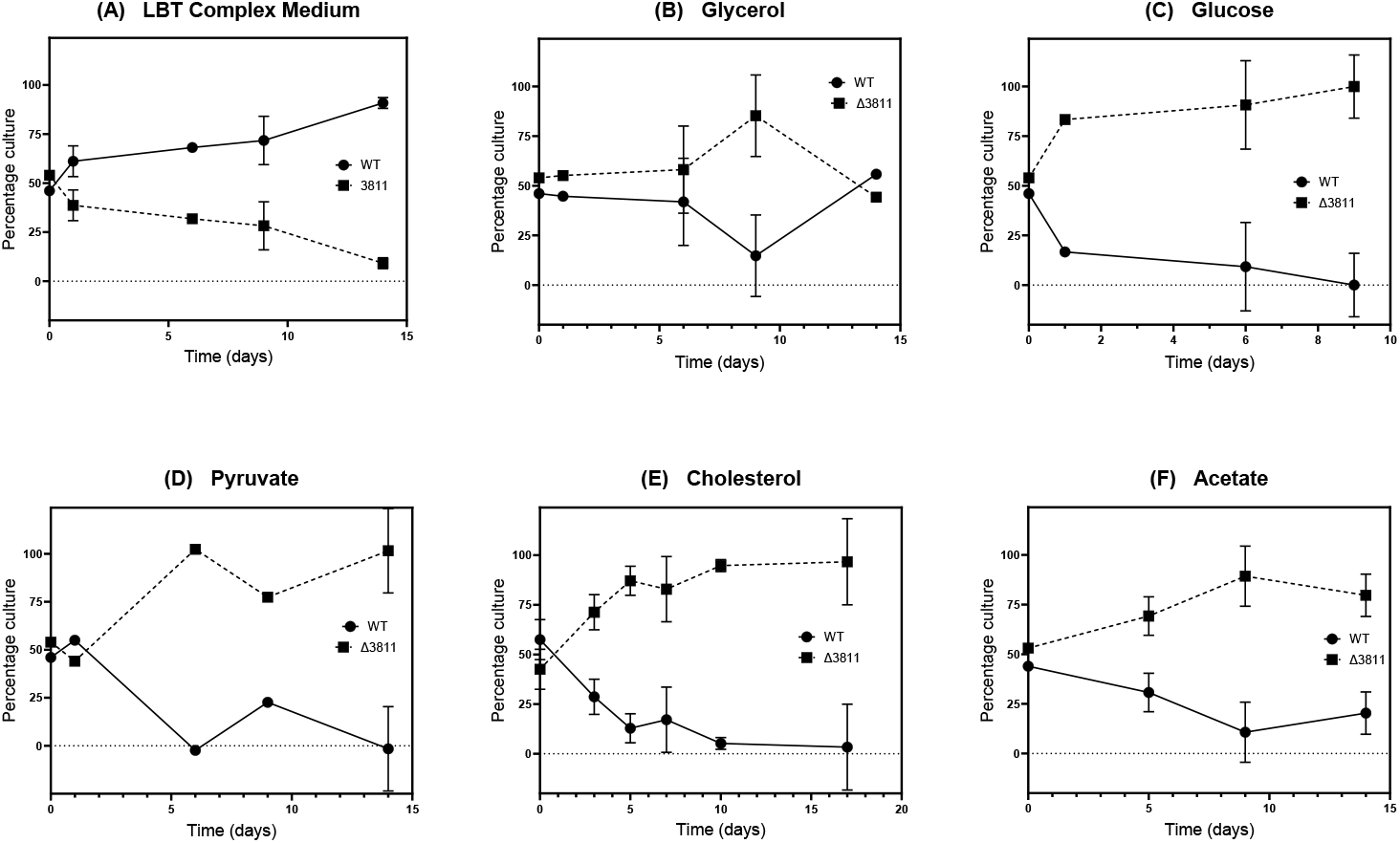
Exponential phase competition experiments. *M. smegmatis* wildtype (WT) and the Δ3811 mutant were grown to mid-exponential phase in HdB medium (OD_600_ = 0.5) with the indicated carbon source and then 50 μl of each culture was inoculated into 5 ml fresh HdB medium incubated and then sub-cultured every day to maintain the mixed culture in exponential phase for the duration of the experiment.

**Figure 2:**
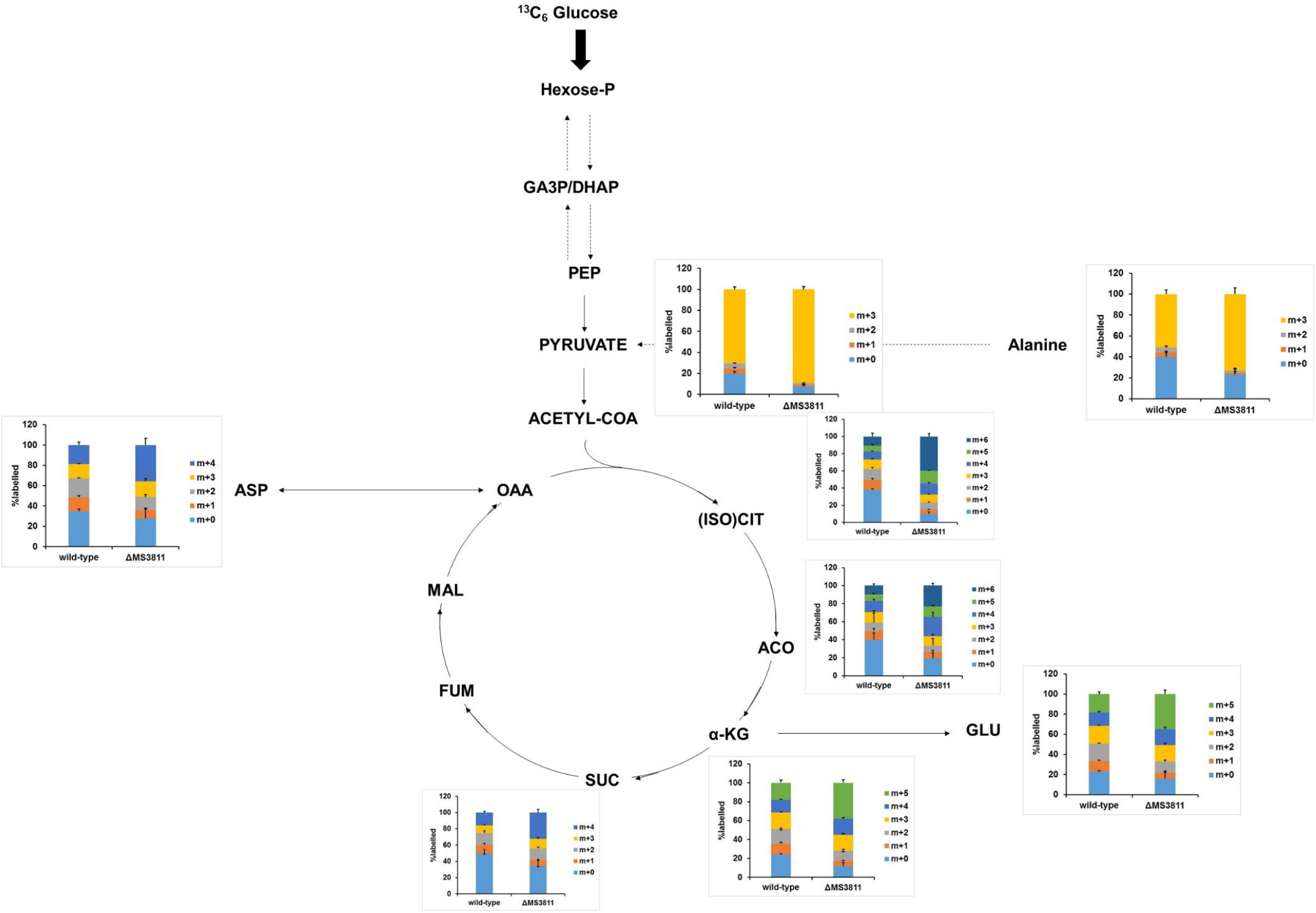
Relative isotolopogue distribution of key metabolites from the central carbon metabolism. Metabolites were extracted as described in the Material ad Methods section at half-doubling time after the labelling with [^13^C_6_-Glucose]. Excess of values were obtained by subtracting the natural ^13^C abundance of 1.11%. The M+0 bar in the diagram represents molecules carrying only ^12^C atom, the M+1 bar stands for molecules carrying only one ^13^C atom (position unknown), the M+2 bar stands for molecules carrying two ^13^C, and so on. Error bars indicate standard deviations from two biological and three technical replicates.

These data support a model whereby MSMEG_3811 acts as a brake on central carbon metabolism, as in the ΔMSEMEG_3811 mutant, there is increased incorporation of carbon consistent with increased carbon flux.

### The mycobacterial single domain USP interacts directly with key central metabolic enzymes

We then reasoned that if the mycobacterial single domain USP regulates flux then it might interact with metabolic enzymes. To test this hypothesis, we used an *in vivo* protein crosslinking approach to look for protein-protein interactions in exponentially growing mycobacteria (33).

For this part of the study and to extend the relevance of this work to *M. tuberculosis* we used the essential single domain USP Rv1636, which is the *M. tuberculosis* homologue of MSMEG_3811. Rv1636 was expressed in an *M. smegmatis groEL1ΔC* strain from the pHEH-N vector, which expresses Rv1636 tagged at its N-terminus with a His-Strep-Strep tag from the mycobacterial hsp60 promoter (33). *M smegmatis groEL1ΔC* was grown alongside a control strain without the vector and the bacteria treated with formaldehyde to stop growth and cross-link *in vivo* protein complexes, which were then purified using a two-step procedure of StrepTactin FPLC followed by nickel affinity purification. Following trypsin digestion proteins complexing with Rv1636 were identified by mass spectrometry.

Twenty-eight proteins with at least 5 unique peptides were detected in the Rv1636 expressing strain but not in the control stain (Table 2). Nine of these were proteins that were also pulled down using a different bait protein (MSMEG_1807, Tuf, DnaK MSMEG_1842,) (33) and/or potentially biotinylated proteins (MSMEG_1813, MSMEG_6391, MSMEG_4717, MSMEG_5404, MSMEG_5086) which will bind StrepTactin with high affinity and may be false positives, and another was MSMEG_3811. Of the remaining 18 proteins the largest group were metabolic enzymes, and included the glycolytic enzymes pyruvate kinase, pyruvate dehydrogenase and pyruvate synthase, as well as glycerol kinase and additional metabolic enzymes including the TCA cycle enzyme aconitate hydratase (aconitase) and glycogen phosphorylase the initiator of glycogen breakdown for carbon entry into glycolysis and 6-phosphgluconate dehydrogenase of the pentose phosphate pathway). These data are consistent with an ability of the conserved single domain USP from mycobacteria to interact directly with key regulated enzymes from the central carbon metabolism.

**Table 2:** Identification of Rv1636 interactor proteins using following in vivo crosslinking. Proteins identified with at least 5 unique peptides following LC-MS are reported.

| UniProt<br>Accession | Gene Name | Description | Unique<br>peptides |
| --- | --- | --- | --- |
| **Information Pathways** |  |  |  |
| **P60281** | *rpoB* (MSMEG_1367) | DNA directed RNA polymerase Beta subunit | 12 |
| **A0QS66** | *rpoC* (MSMEG_1368) | DNA directed RNA polymerase Beta prime subunit | 25 |
| **A0QS98** | *tuf* (MSMEG_1401) | Elongation factor Tu | 25 |
| **A0QVQ5** | *gpsI* (MSMEG_2656) | Guanosine phosphate synthetase I/ polyribonucleotide nucleotidyltransferase | 5 |
| **Virulence, Detoxification, Adaptation** |  |  |  |
| **A0QYW6** | MSMEG_3811 | Universal stress protein family | 8 |
| **A0QQU5** | *groL1* (MSMEG_0880) | 60 kDa chaperonin 1 GroEL | 35 |
| **A0QQC8** | *dnaK* (MSMEG_0709) | Chaperone protein DnaK | 28 |

| Intermediary Metabolism and Respiration |  |  |  |
| --- | --- | --- | --- |
| **A0R0B0** | *aceE* (MSMEG_4323) | Pyruvate dehydrogenase E1 component | 5 |
| **A0R171** | MSMEG_4646 | Pyruvate synthase | 5 |
| **A0R726** | *glpK* (MSMEG_6756) | Glycerol kinase | 14 |
| **A0QXA3** | *pyk* (MSMEG_3227) | Pyruvate kinase | 11 |
| **A0QX20** | *acnA* (MSMEG_3143) | Aconitate hydratase 1 | 8 |
| **A0QYE7** | *gnd*<br>(MSMEG_3632) | 6-phosphogluconate dehydrogenase decarboxylating | 5 |
| **A0R1Y2** | MSMEG_4915 | Glycogen phosphorylase | 6 |
| **A0R2H8** | *pruA*<br>(MSMEG_5119) | 1-pyrroline-5-carboxylate dehydrogenase. Proline metabolism | 7 |
| **A4ZHR8** | *ahcY*<br>(MSMEG_1843) | adenosylhomocysteinase | 17 |
| **A0QY79** | *bfr*<br>(MSMEG_3564) | Bacterioferritin | 5 |
| **A0R3L1** | MSMEG_5512 | Magnesium chelatase | 7 |
| **A0R574** | *clpC1*<br>(MSMEG_6091) | ATP-dependent Clp protease ATP-binding subunit ClpC1 | 28 |
| **Cell wall and cell processes** |  |  |  |
| **A0R1C3** | *ettA*<br>(MSMEG_4700) | ABC-transporter protein, ATP binding component, and likely energy-dependent translational throttle protein EttA | 5 |
| **A0QV32** | *ffh*<br>(MSMEG_2430) | Signal recognition particle protein | 7 |
| **P71533** | *secA1*<br>(MSMEG_1881) | Protein translocase subunit SecA 1 | 5 |
| **Lipid Metabolism** |  |  |  |
| **A0R616** | MSMEG_6391 | Propionyl-CoA carboxylase beta chain <sup>a</sup> | 30 |
| **A0QTE7** | MSMEG_1813 | Propionyl-CoA carboxylase beta chain <sup>a</sup> | 25 |
| **A0R1D9** | MSMEG_4717 | Carboxyl transferase domain protein <sup>a</sup> | 6 |
| **A0QTE1** | MSMEG_1807 | Acetyl-/propionyl-CoA carboxylase alpha chain <sup>a</sup> | 13 |
| **A0R3A6** | MSMEG_5404 | Propionate CoA ligase | 9 |
| **A0R2E8** | MSMEG_5086 | Very long chain acyl CoA synthetase | 6 |

## Discussion

Except for a few bacteria, such as mycoplasma species, universal stress proteins are present in almost all bacteria and archaea. They are broadly associated with bacterial responses to environmental stress (1, 4). The paradigm is UspA from *E. coli*, which was identified as being upregulated in response to a wide variety of stresses and *uspA* mutation resulted in reduced survival under a variety of growth-arresting conditions, while *uspA* overexpression leds to a growth-arrested state (8, 10, 38). *In vitro* studies have shown that *E. coli* UspA can autophosphorylate, but the physiological role of this modification is not known and a tyrosine phosphoprotein TypA is thought to phosphorylate UspA at serine/threonine residues *in vivo* (39, 40).

Most bacteria have multiple paralogs of USP_s_ and individual *usp* gene deletion often does not lead to distinctive phenotype and the consequent inability to use classical suppressor mutant approaches has hindered the development of a biochemical and functional understanding of their cellular roles (1). *M. tuberculosis* possesses 10 USP homologues of which 6 are part of the dormancy or DosR regulon and consequently up-regulated under hypoxia, a stress that leads to entry of this aerobic bacterium into a non-replicating persistent state, thought to resemble that of latent TB infections (7). Knock-out mutants of individual genes encoding the USP_s_ Rv1996, Rv2005c, Rv2026c and Rv2028c showed no long-term survival defect under hypoxia or a range of stress conditions (41). The tandem domain USP Rv2623 is an ATP-binding protein that is hypoxia-induced as part of the DosR regulon but an *rv2623* mutant exhibited a hypervirulent phenotype, while overexpression attenuated growth *in vitro* (20).

*M. tuberculosis* has only one single domain USP, Rv1636, and it is the *M. tuberculosis* USP that is most closely related to the *E. coli* UspA, although Rv1636 differs in having a conserved NTP binding domain, and so is more like the Usp MJ0577 from *Methanococcus janaschi*. The *M. leprae* genome is viewed as having a minimal mycobacterial gene set due to the multiple gene deletions and disruptions that have resulted in its preserving only the most vital genes that are required for its obligate parasitism (42). One of these genes encodes a single domain USP ML1390 that has 89% sequence identity to Rv1636 from *M. tuberculosis*, suggesting that this mycobacterial USP is associated with critical, core cellular functions. While *M. smegmatis* has two single domain USP_s_, MSMEG_3811 is the one most closely related to *M. tuberculosis* Rv1636 and ML1390, with 78.2% and 75.7% sequence identity respectively, and so it is highly probable that these 3 mycobacterial USP_s_ have similar cellular functions.

We aimed to study the function of Rv1636 but were unable to construct a knockout mutant in its gene, a finding that fits with its reported essentially for growth in vitro (36). Therefore, for much of this study we turned to MSMEG_3811 as we were able to knock out its gene and so compare mutant and wild type phenotypes. Initially, this was not particularly informative, as we could not find a stress survival defect in the mutant, despite looking at a very wide range of stresses.. However, quantitative MS measurements determined that MSMEG_3811 protein was present at similar levels in exponential and in stationary phase. This contrasts with the hypoxia induction of most mycobacteria USP_s_ and the growth phase dependence of many others including *E. coli* UspA (1, 4, 7). We therefore set up exponential phase competition experiments between wildtype and mutant which gave striking data. The presence of MSMEG_3811 was conditionally advantageous during *in vitro* growth. It was of benefit and allowed the wildtype to outcompete the mutant in the rich, complex nutritional environment provided by LB medium, but was clearly disadvantageous with a more restricted diet of HdB medium with a single carbon source.

This led us to look at whether the mycobacterial singe domain USP has a role in metabolic control. To test this, we performed ^13^C-stable isotope tracing to investigate whether there was an alteration in the turnover of central metabolites in the parental and the MSMEG_3811 deletion mutant. The data was clear; the absence of MSMEG_3811 led to an increase in both the proportion of label incorporation into the metabolites involved in central carbon metabolism as well as an increase in isotopologues containing a high number of carbons and a decrease in the labelling with isotopologues having a low number.

An *in vivo* crosslinking and pull-down experiment and indicated that Rv1636 interacted directly with central metabolic pathway enzymes including the key regulated glycolytic enzymes pyruvate kinase, pyruvate dehydrogenase and pyruvate synthase, the TCA cycle enzyme aconite hydratase, 6-phosphogluconate dehydrogenase of the pentose phosphate pathway as well as glycogen phosphorylase which catalyses the first step in glycogen degradation by bacteria. These data are consistent with a model whereby this mycobacterial USP regulates the flux through central metabolic pathways.

Many bacteria when presented with multiple carbon sources use catabolite repression as a regulatory mechanism to maximise growth by utilising individual carbon substrates in a favoured sequence and growing with diauxic kinetics. However, *M. tuberculosis* can catabolise multiple carbon sources simultaneously while maintaining monophasic, a process that must require careful control growth (43, 44).

Based on these observations, we can suggest an explanation for the competitive phenotypes whereby in the absence of USP there is a lack of flux control. In a rich medium with multiple carbon sources this may allow the uncontrolled flux of multiple carbon sources through the CMPs and is energetically inefficient and disadvantageous. In contrast when growing *in vitro* on single carbon source the lack of control and the resultant increased flux is advantageous and gives a competitive advantage to the mutant.

The intracellular environment in which a pathogenic mycobacteria finds themselves is nutritionally complex and so (43) careful flux control may be important for optimal growth of the pathogen. We propose the mycobacterial single domain USP proteins can act as dynamic brakes or rheostats on carbon catabolism by regulating flux through central catabolic pathways. Deletion of the USP leads to complete release of the brake and to a loss of control of metabolic flux which is disadvantageous when the bacterium is trying to efficiently coordinate the use of multiple carbon sources, but advantageous under conditions when a single carbon source is available and being used in competition with a wild type strain. As in *M. smegmatis* MSMEG_3811 is constitutively expressed the brake on metabolism must be dynamically controlled in real time and so we propose to call this rheostatic braking and that the USP is acting as a rheostat to regulate metabolic flux. We suggest that the mycobacterial USP_s_ achieves this control by interaction with the key regulatable enzymes, such as pyruvate kinase, pyruvate dehydrogenase and glycerol kinase, that Rv1636 interacted with in the *in vivo* crosslinking experiments. Rv1636 and MSMEG_3811 have been shown to bind cAMP and it is an intriguing possibility, given that cAMP is the regulator of catabolite repression that coordinates the ordered sequential use of carbon sources and diauxic growth, that it has a role in mycobacteria to coordinate carbon flux (23).

There are examples of USP_s_ having metabolic roles in other bacteria. In the photosynthetic bacterium *Rhodobacter sphaeroides* UspA was found to interact with the cytochrome *bc1* complex of the respiratory chain resulting in an increase in *bc1* complex activity and a decrease in superoxide generation (16). The *M. tuberculosis* USP Rv2623 interacts with a putative ATP binding cassette (ABC) transporter Rv1747, negatively regulating the activity of this putative transporter of phosphatidyl-myo-inositol mannosides (PIMs) (22), while overexpression of the *M. tuberculosis* USP Rv2624c in *M. smegmatis* increased the intracellular arginine levels (45).

There is support for *E. coli* UspA being involved in carbon metabolism. A *ΔuspA* mutant had a lag in starting growth in minimal medium with a variety of carbon sources, and when it entered exponential growth it showed a diauxic growth phenotype with glucose or gluconate (9). Particularly interesting and relevant to the data presented in this paper, was that the mutant utilised glucose more rapidly than the wildtype and produced more acetate suggesting a disturbance to carbon metabolism (9). Furthermore, the *uspA* gene was regulated by fructose-6-phospahte levels suggesting its regulation is influenced by metabolite levels (46). These data suggest that *E. coli* UspA might also act as a metabolic rheostat and this should be directly investigated.

The data presented in this paper are an important contribution to developing an understanding of the biochemical roles of USP_s_ as well as their specific roles in mycobacteria including *M. tuberculosis*. The essentiality Rv1636 together with ML1390 being the sole USP in the *M. leprae* genome lends support to these proteins playing important roles in infection. While *M. tuberculosis* will grow in the phagosome of macrophages it can also escape this compartment to survive extracellularly and within the environment of granulomas. During infection *M. tuberculosis* is in a complex and varied nutritional environmental and flux control making use of USP-mediated rheostatic braking is potentially an important mechanism to optimise catabolism.

## Materials and Methods

### Bacterial Strains and Growth Media

*M. smegmatis* mc^2^155 was routinely cultured at 37°C in LB medium (0.5% (w/v) NaCl, 0.5% (w/v) yeast extract, 1.0% (w/v) tryptone) supplemented with 0.05% (w/v) Tween 80. For carbon starvation and hypoxic growth experiments, the cells were grown in Hartmans-de Bont (HdB) medium as described previously (24–26). For the heat shock experiments, cultures were grown into hypoxic stationary phase, and the cultures were transferred to a 55 °C incubator at 48 h postinoculation. For culture on solid medium, we added 2% (w/v) agar to medium prior to autoclaving. Bacterial viability was determined by plating 20 μl serial dilutions onto solid media, and growth was enumerated after 3 days.

### Construction of the MSMEG_3811 Mutant and Complementation

Mutant *M. smegmatis* strains were obtained following an established protocol (27). Briefly, a linear construct was generated so that it contained a hygromycin cassette flanked by 500 bp homologous to the sequence found up- and downstream of MSMEG_3811 in *M. smegmatis*. The construct was amplified by PCR and electroporated into *M. smegmatis* mc2155 cells harbouring the pJV53 plasmid and, following recovery, plated onto 7H10 medium containing hygromycin (20 μg/ml). Genomic DNA was purified from potential mutants and assessed for the presence of the hygromycin cassette as well as its insertion into the correct genetic locus by PCR.

### Competition Experiments

*M. smegmatis* mc^2^155 and the MSMEG_3811 mutant strains to be competed were grown in HdB medium with the specific carbon source to be tested to an OD_600_ of 0.5 and then 50 μl of each culture mixed into 5 ml of fresh HdB, grown at 37°C and sub-cultured every day by transferring 100 μl of the competition mixture into fresh HdB medium to keep the bacteria in exponential phase for the duration of the experiment. Samples were taken for mutant and total viable counting by plating onto LB with or without hygromycin, respectively. The following carbon sources were added to HdB for the various competition experiments: glycerol (0.2 or 0.4 % v/v), glucose (0.2% w/v), cholesterol (0.2% w/v), propionate (0.2% w/v), acetate (0.2% w/v).

### Multiple Reaction Monitoring Mass spectrometry (MRM-MS)

Signature peptides corresponding to USP 3811 (MSMEG_3811) and S2 30S subunit ribosomal protein (MSMEG_2519) were selected for synthesis based on their length, uniqueness and signal intensity of MRM transitions. For MSMEG_3811 the peptide was AGQIAAASNA and the transitions were 2y6, 2y7 and 2y8 and for the 30S ribosomal subunit S2 (MSMEG_2519) the peptide was VIASAVAEGLQAR and the transitions 2y6, 2y7 and 2y8 and 2y10. The lyophilised internal standard peptides (synthesised by JPT Peptide Technologies GmbH) were suspended in 80% 0.1 M NH_4_HCO_3_ and 20% acetonitrile to 5 μM concentration. 25 μL of urea solubilised lysates of *M. smegmatis* were mixed with 0.5 μl of MSMEG_3811 AGQIAAASNA peptide and 1 μL of S2 peptide VIASAVAEGLQAR. The samples were diluted with 225 μL of 57.6 mM NH_4_HCO_3_ containing 1 mM TCEP and the mixtures treated with 1 μg of sequencing grade modified trypsin (Promega), incubated at 37 °C overnight and then acidified by addition of formic acid to 0.1 % v/v and stored at −20 °C. Three biological replicates of *M. smegmatis* lysates were analysed by MRM-MS as described in (28). MRM analysis generated spectral peaks for light peptide transitions from lysates and heavy peptide transitions from the internal peptide standards. The peaks were integrated using Skyline software (https://www.skylinesoft.com/) and signals from light transitions divided by those from heavy transitions. The resulting ratios for different transitions were averaged for each specific peptide and the signal normalised to 1 mg/mL total protein in lysates.

### Stable isotope tracing samples preparation

For targeted metabolomics profiling, strains were cultured as described previously. Briefly, mycobacteria were grown initially in HdB liquid medium containing the carbon sources of interest until the OD_600_ reached ~0.8-1. Bacteria were then inoculated onto 0.22 μm nitrocellulose filters under vacuum filtration. Mycobacterial-laden filters were then placed on top of chemically equivalent agar media (described above) and allowed to grow at 37°C for 5 doubling times to generate enough biomass for targeted metabolomics studies. Filters were then transferred into HdB agar plates supplemented with 0.2% [U-^13^C_6_] glucose for a half-doubling time. Bacteria were then metabolically quenched by plunging the filters into the extraction solution composed of acetonitrile/methanol/H_2_O (2:2:1). Small molecules were extracted by mechanical lysis of the entire bacteria-containing solution with 0.1 mm acid-washed zirconia beads for 1 min using a FastPrep (MPBio^®^) set at 6.0 m/s. Lysates were filtered through 0.22 μm Spin-X column filters (Costar^®^). Bacterial biomass of individual samples was determined by measuring the residual protein content of the metabolite extracts using the BCA assay kit (Thermo^®^) (29–32).

### Liquid-chromatography-mass spectrometry

Aqueous normal phase liquid chromatography was performed using an Agilent 1290 Infinity II LC system equipped with a binary pump, temperature-controlled auto-sampler (set at 4°C) and temperature-controlled column compartment (set at 25°C) containing a Cogent Diamond Hydride Type C silica column (150 mm × 2.1 mm; dead volume 315 μl). A flow rate of 0.4 ml/min was used. Elution of polar metabolites was carried out using solvent A consisting of deionized water (resistivity ~18 MΩ cm) and 0.2% acetic acid and solvent B consisting of 0.2% acetic acid in acetonitrile. The following gradient was used: 0 min 85% B; 0-2 min 85% B; 3-5 min to 80% B; 6-7 min 75% B; 8-9 min 70% B; 10-11 min 50% B; 11.1-14 min 20% B; 14.1-25 min hold 20% B followed by a 5 min re-equilibration period at 85% B at a flow rate of 0.4 ml/min. Accurate mass spectrometry was carried out using an Agilent Accurate Mass 6545 QTOF apparatus. Dynamic mass axis calibration was achieved by continuous infusion, post-chromatography, of a reference mass solution using an isocratic pump connected to an ESI ionization source operated in the positive-ion mode. The nozzle voltage and fragmentor voltage were set at 2,000 V and 100 V, respectively. The nebulizer pressure was set at 50 psig, and the nitrogen drying gas flow rate was set at 5 l/min. The drying gas temperature was maintained at 300°C. The MS acquisition rate was 1.5 spectra/sec, and *m/z* data ranging from 50-1,200 were stored. This instrument enabled accurate mass spectral measurements with an error of less than 5 parts-per-million (ppm), mass resolution ranging from 10,000-45,000 over the *m/z* range of 121-955 atomic mass units, and a 100,000-fold dynamic range with picomolar sensitivity. The data were collected in the centroid 4 GHz (extended dynamic range) mode. Detected *m/z* were deemed to be identified metabolites based on unique accurate mass-retention time and MS/MS fragmentation identifiers for masses exhibiting the expected distribution of accompanying isotopomers. Typical variation in abundance for most of the metabolites remained between 5 and 10% under these experimental conditions. Under the experimental conditions described above using [U-^13^C_6_] glucose (99%), the extent of ^13^C labelling for each metabolite was determined by dividing the summed peak height ion intensities of isotopologue ^13^C-labelled species by the ion intensity of both labelled and unlabelled species using the software Agilent Profinder version B.8.0.00 service pack 3. Data are presented as the mean ± standard error of the mean from 2 biological replicates and 3 technical replicates per condition. Unpaired two-tailed Student’s *t*-tests were used to compare values, with p < 0.05 considered significant

### In vivo crosslinking to identify interacting partner proteins of the single domain USP Rv1636

This was done as described previously using the pHEH-N (Rv1636) construct exprressing the Mtb single domnain USP Rv1636, which expresses Rv1636 tagged at its N-terminus with a His-Strep-Strep tag from the mycobacterial hsp60 promoter (33). Briefly, *M. smegmatis groEL1ΔC* (34) was trasnformed with pHEH (Rv1636) and grown in LB + 0.05% Tween 80 for 24h and complexes crosslinked with 0.4% formaldehyde. Formaldehyde cross-linked protein complexes were purified from cell extracts by sequential purification on StrepTactin and nickel affinity columns. The crosslinks were reversed by heating at 95°C for 20 mins and the proteins separated by SDS-PAGE (see Figure S X). No proteins were purified from the control cultures wheras a number of bands co—purified with Rv1636. The protein presnet in the bands were identified following trypsin digestion and LC-MS as described previously (Table 2) (33).

## Acknowledgments

This work was funded in part by the Wellcome Trust (086087/Z/08/Z).

## Supplementary Figure

**Fig. S1.**
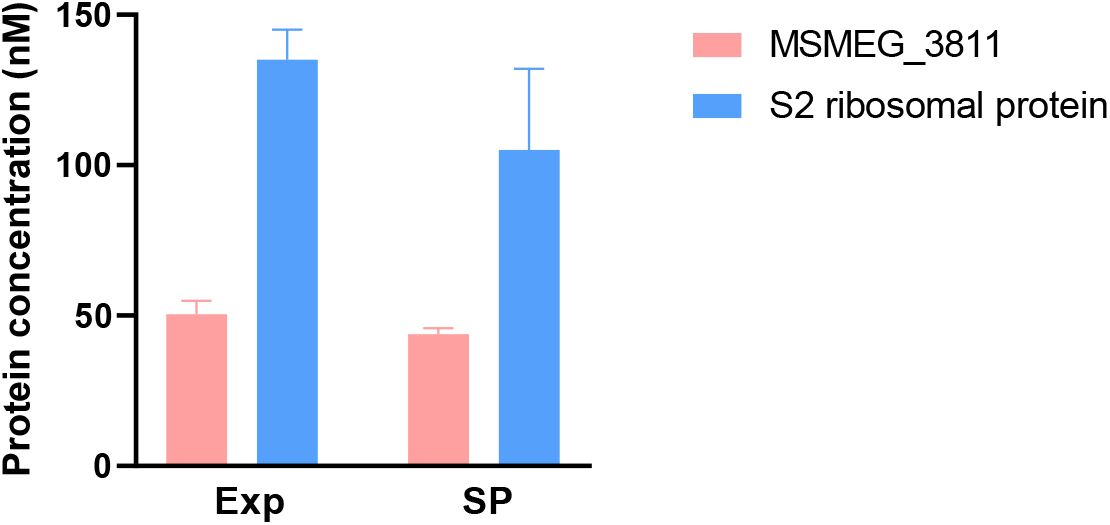
MRM-MS analysis of MSMEG-3811 protein concentrations in *M. smegmatis*. The expression levels of MSMEG_3811 and the ribosomal protein S2 were analysed in exponentially growing and stationary phase *M. smegmatis*. There was no significant differences in the concentrations of these proteins between exponential and stationary phase calculated using a One-Way ANOVA with Tukey multiple pairwisecomparisons between means.

## References

1. A. C. Vollmer, S. J. Bark, Twenty-Five Years of Investigating the Universal Stress Protein: Function, Structure, and Applications. Adv Appl Microbiol 102, 1–36 (2018).

2. D. A. Siegele, Universal stress proteins in Escherichia coli. J Bacteriol 187, 6253–6254 (2005).

3. K. L. Tkaczuk et al., Structural and functional insight into the universal stress protein family. Evolutionary applications 6, 434–449 (2013).

4. K. Kvint, L. Nachin, A. Diez, T. Nystrom, The bacterial universal stress protein: function and regulation. Curr Opin Microbiol 6, 140–145 (2003).

5. T. I. Zarembinski et al., Structure-based assignment of the biochemical function of a hypothetical protein: a test case of structural genomics. Proc Natl Acad Sci U S A 95, 15189–15193 (1998).

6. M. C. Sousa, D. B. McKay, Structure of the universal stress protein of Haemophilus influenzae. Structure 9, 1135–1141 (2001).

7. R. O’Toole, H. D. Williams, Universal stress proteins and Mycobacterium tuberculosis. Res Microbiol 154, 387–392 (2003).

8. T. Nystrom, F. C. Neidhardt, Cloning, mapping and nucleotide sequencing of a gene encoding a universal stress protein in Escherichia coli. Mol Microbiol 6, 3187–3198 (1992).

9. T. Nystrom, F. C. Neidhardt, Isolation and Properties of a Mutant of Escherichia-Coli with an Insertional Inactivation of the uspA Gene, Which Encodes a Universal Stress Protein. Journal of Bacteriology 175, 3949–3956 (1993).

10. T. Nystrom, F. C. Neidhardt, Expression and role of the universal stress protein, UspA, of Escherichia coli during growth arrest. Mol Microbiol 11, 537–544 (1994).

11. W. T. Liu et al., Role of the universal stress protein UspA of Salmonella in growth arrest, stress and virulence. Microb Pathog 42, 2–10 (2007).

12. N. Boes, K. Schreiber, E. Hartig, L. Jaensch, M. Schobert, The Pseudomonas aeruginosa universal stress protein PA4352 is essential for surviving anaerobic energy stress. J Bacteriol 188, 6529–6538 (2006).

13. K. Schreiber et al., Anaerobic survival of Pseudomonas aeruginosa by pyruvate fermentation requires an Usp-type stress protein. J Bacteriol 188, 659–668 (2006).

14. R. C. Fink et al., FNR is a global regulator of virulence and anaerobic metabolism in Salmonella enterica serovar Typhimurium (ATCC 14028s). J Bacteriol 189, 2262–2273 (2007).

15. W. Chen, K. Honma, A. Sharma, H. K. Kuramitsu, A universal stress protein of Porphyromonas gingivalis is involved in stress responses and biofilm formation. FEMS Microbiol Lett 264, 15–21 (2006).

16. T. Su, Q. Wang, L. Yu, C. A. Yu, Universal Stress Protein Regulates Electron Transfer and Superoxide Generation Activities of the Cytochrome bc1 Complex from Rhodobacter sphaeroides. Biochemistry 54, 7313–7319 (2015).

17. M. Kundu, J. Basu, Applications of Transcriptomics and Proteomics for Understanding Dormancy and Resuscitation in Mycobacterium tuberculosis. Front Microbiol 12, 642487 (2021).

18. R. Heermann et al., The universal stress protein UspC scaffolds the KdpD/KdpE signaling cascade of Escherichia coli under salt stress. J Mol Biol 386, 134–148 (2009).

19. H. D. Park et al., Rv3133c/dosR is a transcription factor that mediates the hypoxic response of Mycobacterium tuberculosis. Mol Microbiol 48, 833–843 (2003).

20. J. E. Drumm et al., Mycobacterium tuberculosis Universal Stress Protein Rv2623 Regulates Bacillary Growth by ATP-Binding: Requirement for Establishing Chronic Persistent Infection. Plos Pathogens 5, - (2009).

21. S. M. Hingley-Wilson, K. E. Lougheed, K. Ferguson, S. Leiva, H. D. Williams, Individual Mycobacterium tuberculosis universal stress protein homologues are dispensable in vitro. Tuberculosis (Edinb) 90, 236–244 (2010).

22. L. N. Glass et al., Mycobacterium tuberculosis universal stress protein Rv2623 interacts with the putative ATP binding cassette (ABC) transporter Rv1747 to regulate mycobacterial growth. PLoS Pathog 13, e1006515 (2017).

23. A. Banerjee et al., A universal stress protein (USP) in mycobacteria binds cAMP. J Biol Chem 290, 12731–12743 (2015).

24. R. O’Toole et al., A two-component regulator of universal stress protein expression and adaptation to oxygen starvation in Mycobacterium smegmatis. J Bacteriol 185, 1543–1554 (2003).

25. M. J. Smeulders, J. Keer, R. A. Speight, H. D. Williams, Adaptation of Mycobacterium smegmatis to stationary phase. J Bacteriol 181, 270–283 (1999).

26. J. Keer, M. J. Smeulders, H. D. Williams, A purF mutant of Mycobacterium smegmatis has impaired survival during oxygen-starved stationary phase. Microbiology 147, 473–481 (2001).

27. J. C. van Kessel, G. F. Hatfull, Recombineering in Mycobacterium tuberculosis. Nat Methods 4, 147–152 (2007).

28. J. Schumacher et al., Nitrogen and carbon status are integrated at the transcriptional level by the nitrogen regulator NtrC in vivo. MBio 4, e00881–00813 (2013).

29. S. Rebollo-Ramirez, G. Larrouy-Maumus, NaCl triggers the CRP-dependent increase of cAMP in Mycobacterium tuberculosis. Tuberculosis (Edinb) 116, 8–16 (2019).

30. G. Larrouy-Maumus et al., Cell-Envelope Remodeling as a Determinant of Phenotypic Antibacterial Tolerance in Mycobacterium tuberculosis. ACS Infect Dis 2, 352–360 (2016).

31. A. Gouzy et al., Mycobacterium tuberculosis nitrogen assimilation and host colonization require aspartate. Nat Chem Biol 9, 674–676 (2013).

32. G. Larrouy-Maumus et al., Discovery of a glycerol 3-phosphate phosphatase reveals glycerophospholipid polar head recycling in Mycobacterium tuberculosis. Proc Natl Acad Sci U S A 110, 11320–11325 (2013).

33. K. E. Lougheed, M. H. Bennett, H. D. Williams, An in vivo crosslinking system for identifying mycobacterial protein-protein interactions. J Microbiol Methods 105, 67–71 (2014).

34. E. E. Noens et al., Improved mycobacterial protein production using a Mycobacterium smegmatis groEL1DeltaC expression strain. BMC Biotechnol 11, 27 (2011).

35. V. Pelicic, J. M. Reyrat, B. Gicquel, Generation of unmarked directed mutations in mycobacteria, using sucrose counter-selectable suicide vectors. Mol Microbiol 20, 919–925 (1996).

36. J. E. Griffin et al., High-resolution phenotypic profiling defines genes essential for mycobacterial growth and cholesterol catabolism. PLoS Pathog 7, e1002251 (2011).

37. J. E. Drumm et al., Mycobacterium tuberculosis universal stress protein Rv2623 regulates bacillary growth by ATP-Binding: requirement for establishing chronic persistent infection. PLoS Pathog 5, e1000460 (2009).

38. T. Nystrom, F. C. Neidhardt, Isolation and properties of a mutant of Escherichia coli with an insertional inactivation of the uspA gene, which encodes a universal stress protein. J Bacteriol 175, 3949–3956 (1993).

39. P. Freestone, T. Nystrom, M. Trinei, V. Norris, The universal stress protein, UspA, of Escherichia coli is phosphorylated in response to stasis. J Mol Biol 274, 318–324 (1997).

40. P. Freestone, M. Trinei, S. C. Clarke, T. Nystrom, V. Norris, Tyrosine phosphorylation in Escherichia coli. J Mol Biol 279, 1045–1051 (1998).

41. S. Hingley-Wilson, K. Lougheed, K. Ferguson, S. Leiva, H. Williams, Individual Mycobacterium tuberculosis universal stress protein homologues are dispensable in vitro. Tuberculosis 90, 236–244 (2010).

42. S. T. Cole et al., Massive gene decay in the leprosy bacillus. Nature 409, 1007–1011 (2001).

43. K. Y. Rhee et al., Central carbon metabolism in Mycobacterium tuberculosis: an unexpected frontier. Trends Microbiol 19, 307–314 (2011).

44. L. P. de Carvalho et al., Metabolomics of Mycobacterium tuberculosis reveals compartmentalized co-catabolism of carbon substrates. Chem Biol 17, 1122–1131 (2010).

45. Q. Jia et al., Universal stress protein Rv2624c alters abundance of arginine and enhances intracellular survival by ATP binding in mycobacteria. Sci Rep 6, 35462 (2016).

46. O. Persson, A. Valadi, T. Nystrom, A. Farewell, Metabolic control of the Escherichia coli universal stress protein response through fructose-6-phosphate. Mol Microbiol 65, 968–978 (2007).

